# Fine-tuned synthetic transcription factors for production of 3′-phosphoadenosine-5′- phosphosulfate in yeast

**DOI:** 10.64898/2026.02.02.703094

**Authors:** Madhushruti Borah, Shanna Gu, Essa M. Saied, Christoph Arenz, Mattheos Koffas, Gita Naseri

## Abstract

Technologies developed over the past decade have made *Saccharomyces cerevisiae* a promising platform for producing various natural products. Balancing multi-enzyme expression, while maintaining robust microbial growth, remains a limiting factor for engineering long biosynthetic pathways in yeast. Here, we improved the transcriptional capacity of our previously developed isopropyl β-D-1-thiogalactopyranoside (IPTG)-inducible synthetic transcription factors (synTFs) derived from the plant JUB1 DNA-binding domain. To this end, at cysteine positions within surface-exposed loop regions of a JUB1-derived DNA-binding scaffold, we introduced a short peptide to enhance loop flexibility while providing local stability and orientation. The generated synTFs, so-called JUB1-X synTFs, varying in strength, have been successfully used to improve the synthesis of 3’-phosphoadenosine 5’-phosphosulfate (PAPS), a universal sulfate donor necessary for the synthesis of bioactive molecules, including therapeutic glycosaminoglycans and sulfolipids. Using only this engineered yeast strain in simple batch culture, PAPS accumulation of 21.4 ± 5.8 mg g^−1^ cdw was achieved after only 5 hours of inducing the expression of JUB1-X synTFs. Beyond PAPS production, the design principle demonstrated here provides a generalizable strategy to fine-tune other plant-derived synTFs, expanding the regulatory capabilities of existing synTF collections. Together, this work offers a modular, scalable approach to constructing high-performance gene circuits and supports the development of yeast cell factories for complex metabolic and synthetic biology applications.

## INTRODUCTION

The well-developed genetics, solid knowledge of endogenous metabolism, and robustness in large-scale fermentations make traditional baker’s yeast *Saccharomyces cerevisiae* a suitable cell factory for metabolic engineering^1^ As such, it has been broadly used to produce diverse products, e.g., alcohols^2^, hydrocarbons^3^, and proteins^4^. Extending this capability to the microbial production of activated cofactors would further expand the utility of yeast as a platform for complex biochemicals.

Balancing enzyme expression, correct folding, and cofactor supply while maintaining robust microbial growth remains a limiting factor for large-scale productivity. Strong promoters of yeast genes are repetitively implemented in yeast *S. cerevisiae* for metabolic engineering purposes^5^. They may create a metabolic burden on the cell (thus negatively affecting growth), and cells must allocate limited metabolic resources to heterologous metabolite production during the biomass accumulation phase^6^. To mitigate this trade-off, dynamic pathway engineering emerged as an effective strategy to address metabolic stress and cytotoxicity arising from natural product accumulation^6^. Previously developed inducible synthetic transcription factors (synTFs) have been limited by relatively low transcriptional activation, making them suboptimal for high-performance synthetic biology applications^7^. To overcome these limitations, we previously established a new class of chemically inducible synTFs for gene expression in *S. cerevisiae*. To this end, different combinations of plant TF families and activation domains (ADs) derived from plants, viruses, and yeast are being used in yeast. The isopropyl β-D-1-thiogalactopyranoside (IPTG)-inducible *GAL1* promoter (harboring *lacO* site) controls the expression of plant-derived ATFs in yeast. Upon synTF expression, synTF targets synthetic promoters that harbor one or more copies of its binding site upstream of a yeast *CYC1* minimal promoter to drive target gene expression. Among them, JUB1 (JUNGBRUNNEN1)-derived synTF exhibited strong inducible activation and low basal activity, making it an optimal candidate for synTF engineering for enhancing its transcriptional capacity in yeast. In this study, we engineered next-generation JUB1-based synTFs by rationally modifying conserved regions of the NAC DNA-binding domain (DBD), which is crucial for DNA binding and dimerization. We showed that inserting short, stabilized, flexible sequences into structurally sensitive but non-disruptive regions of DBD could improve transcriptional performance of synTFs without altering DNA-binding specificity, while minimizing growth defects compared to commonly used yeast promoters.

As a proof-of-concept, we implemented our JUB1-X synTFs to enhance production of 3′-phosphoadenosine 5′-phosphosulfate (PAPS), the universal sulfate donor required for sulfotransferase-catalyzed reactions for a wide range of biomolecules^8^, including glycosaminoglycans^9^, glycolipids^10^, and small-molecule metabolites^10^. Sulfated glycosaminoglycans, such as chondroitin sulfate, heparin and heparan sulfate, are of important biomedical significance because of their anticoagulant, anti-inflammatory, and antiviral activities^11^. Growing the need for pharmaceuticals for these compounds has promoted interest in sustainable biosynthetic routes, as traditional extraction from animal-derived sources faces challenges such as pathogen contamination, batch-to-batch variability, and supply chain limits^11^. Consequently, insufficient PAPS supply can severely constrain the productivity of engineered sulfation pathways. PAPS biosynthesis involves the sequential action of ATP sulfurylase (ATPS), adenosine 5′-phosphosulfate kinase (APSK), and inorganic pyrophosphatase (PPA)^12, 13^. Enzymatic PAPS synthesis is strongly limited by the cost of ATP, which alone accounts for more than 60% of substrate expenses^14^. In a recent study, Le *et al* (2025) showed that combining ATP regeneration with biotin–streptavidin-mediated enzyme immobilization in yeast reduced ATP consumption, resulting in an improved PAPS titer of 12.02 g/L^15^. However, the complex enzyme immobilization procedures and cofactor recycling systems required in their study limit the approach’s accessibility and scalability. Hence, in this study, we used JUB1-X synthetic transcription factors to induce expression of the key PAPS biosynthetic enzymes ATPS, PPA, and APSK after yeast growth was complete. We showed that using only our growth-decoupled strategy, mediated by JUB1-X synTFs, enables PAPS accumulation of PAPS to 21.4 ± 5.8 mg g^−1^ cdw after 5 h of induction. The approach is simple, scalable, and readily extendable to other synthetic biology and metabolic engineering applications requiring dynamic gene control.

## MATERIALS AND METHODS

### General

Plasmids were constructed by NEBuilder HiFi DNA assembly (New England Biolabs, Frankfurt am Main, Germany) ^16^. The strains used in this study are listed in **Table 1**. Primer and plasmid sequences are given in **Supplementary Table S1** and **Supplementary Table S2**, respectively. PCR amplifications were done using Q5 DNA Polymerase (New England Biolabs, Frankfurt am Main, Germany), or PrimeSTAR GXL DNA Polymerase (Takara Bio, Saint-Germain-en-Laye, France). Amplified DNA parts were gel-purified prior to Sanger sequencing. All primers and oligonucleotides were ordered from IDT (Integrated DNA Technologies Inc., Dessau-Rosslau, Germany). All constructs were confirmed by sequencing (Microsynth Seqlab, Goettingen, Germany).

**Table 1.**
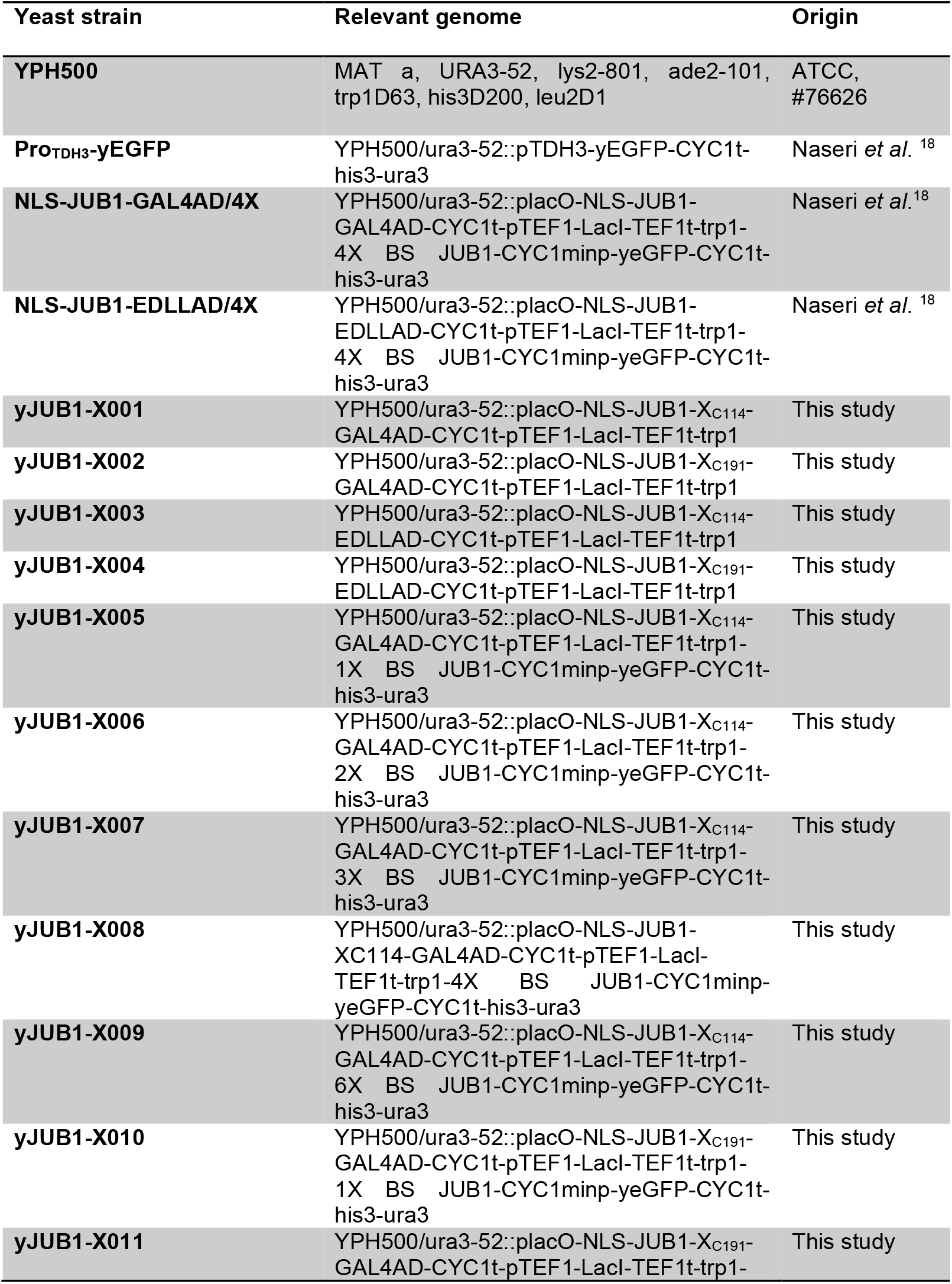
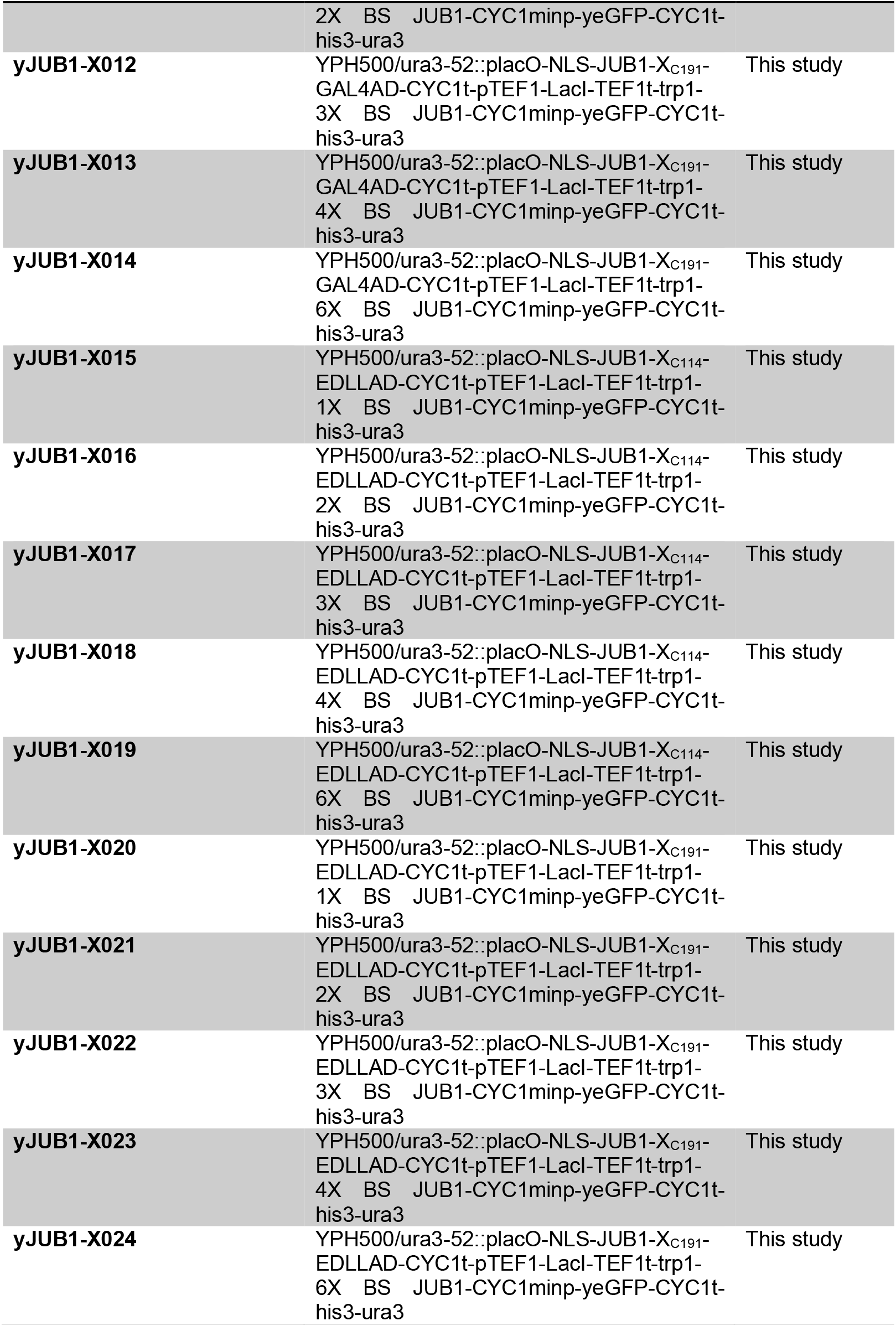
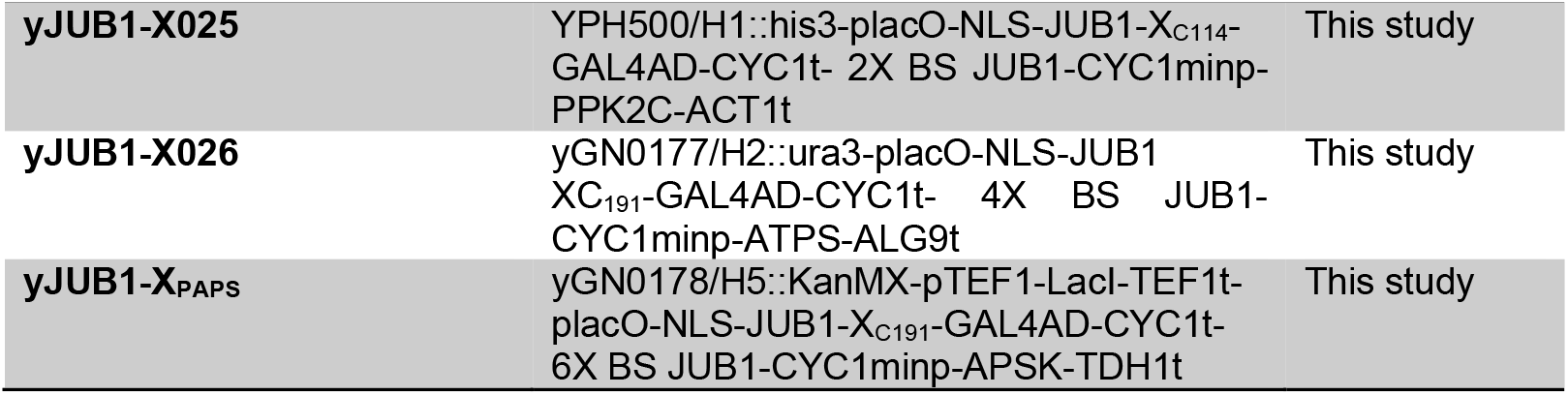
List of parental *S. cerevisiae* strains used in this study.

Plasmids were transformed into *Escherichia coli* NEB 5α (New England Biolabs). Strains were grown in Luria-Bertani (LB) medium with an appropriate *E. coli* selection marker at 37°C. Competent yeast cells and genetic transformation with plasmids or linearized DNA fragments were performed using the LiAc/SS carrier DNA/PEG method^17^. All strains were grown at 30°C in yeast extract peptone dextrose adenine (YPDA)-rich medium or in appropriate synthetic complete (SC) media. Then, verification of the transformation was done using colony PCR followed by sequencing.

### Construction of JUB1-X synTFs Expression Clones

JUB1-X synTF expression clones were constructed based on previously published plasmids pFM003B-JUB1(NLS-JUB1-GAL4AD) and pGN003B-JUB1(NLS-JUB1-EDLLAD), encoding NLS–JUB1 fused to the GAL4 activation domain or the EDLL activation domain^18^. Short peptide insertions (YVG and GER) were added into the JUB1 coding sequence using PCR-based site-directed mutagenesis. To this end, the hexapeptide YVGGER was inserted at positions C114 or C191 using primers SM001/SM002 and SM003/SM004, respectively. The same insertion strategy was applied to both GAL4AD- and EDLLAD-based constructs. Following sequence verification, plasmids pSM01-04 were linearized with *PmeI*, which cuts within the *ura3-52* homology region, and transformed into the yeast strain YPH500. Targeted genomic integration occurred at the *ura3–52* locus via homologous recombination. Positive transformants were selected on SD–Trp medium and confirmed by colony PCR with primers SM005 and SM006, followed by sequencing.

### Construction of JUB1-X synTFs Reporter Clones

Plasmids pSM05,pSM06, pSM07, and pSM08 harbor one, two, three, and four copies of JUB1 binding sites^19^. Six copies of the JUB1 binding site were amplified from pSM09 (synthesized by BioCat, Germany). Next, the reporter plasmid pSM010 harboring six copies of the JUB1 binding site reporter construct was generated by ligating an *EcoR*I/*Xho*I-digested fragment derived from pSM09 into the *EcoR*I/*Xho*I-digested pSM08 using T4 DNA ligase. All reporter plasmids were linearized with *Aat*II and transformed into the yeast strain YPH500 carrying the corresponding JUB1-X synTF expression cassette. Transformants were selected on SD–Ura medium. Correct genomic integration was verified by colony PCR using primers SM006/SM007, followed by Sanger sequencing.

### ATF characterization, flow cytometry, and data analysis

Yeast strains carrying yEGFP reporter constructs were used to assess the transcriptional activity of JUB1-X synTFs under IPTG-inducing conditions. The set included wild-type (WT) strains, a strain with GFP under the constitutive *TDH3* promoter, strains expressing unmodified JUB1 synTFs, and strains expressing JUB1-X variants carrying mutations, and strains expressing JUB1-X variants in combination with various reporters (strains yJUB1-X001 - yJUB1-X025). Single colonies were grown overnight (∼16 h) in selective SC medium at 30°C with 220 rpm. Main cultures were started at an OD600 of ∼0.1 in YPDA medium. Inducing cultures contained 2% (w/v) galactose and 20 mM IPTG, and non-inducing cultures contained 2% (w/v) glucose. Cultures were grown for 4–6 h at 30°C with 220 rpm. Cells were harvested by centrifugation, washed once with phosphate-buffered saline (PBS), and analyzed by flow cytometry. yEGFP fluorescence was measured using a Cytek Aurora CS cytometer, with a minimum of 10,000 cells per sample. The geometric mean of yEGFP fluorescence per cell was calculated in FlowJo.

### Construction of plasmids and donors for PAPS pathway integration

For the PAPS production pathway coupled with ATP generation, the genes PPK2_CpaN_ and ATPS were adopted from *Pseudomonas aeruginosa* and *Kluyveromyces lactis*, respectively, and codon-optimized for *Saccharomyces cerevisiae*, together with APSK from *S. cerevisiae*. The genes PPK2_CpaN_, ATPS, and APSK were expressed under the control of the inducible ATFs NLS-JUB1-XC114-GAL4AD/2X, NLS-JUB1-XC191-GAL4AD/4X, and NLS-JUB1-XC191-GAL4AD/6X, respectively. To construct plasmid pSM11, expression cassette for NLS-JUB1-X_C114_-GAL4AD synTF (primers SM008/SM009, on pSM01), six copies of JUB1 binding sites upstream of the *CYC1* minimal promoter (primers SM010/SM011, on pSM10), oligo fragment encoding coding DNA sequence (CDS) of *PPK2*_*CpaN*_ codon optimized for yeast (synthesized by GenScript, Netherlands), and *ACT1* terminator (primers SM012/SM013, on genomic DNA of *S. cerevisiae* YPH500 strain) were cloned into *Nco*I-digested yeast vector for integration into *H1* locus, harboring yeast selection auxotrphic marker HIS3^20^. To construct plasmid pSM12, expression cassette for NLS-JUB1-X_C191_-GAL4AD synTF (primers SM014/SM009, on pSM02), four copies of JUB1 binding sites upstream of the *CYC1* minimal promoter (primers SM010/SM011, on pSM008), *ATPS* CDS (primers SM0015/SM016, on genomic DNA of *S. cerevisiae* YPH500 strain) and *ALG9* terminator (primers SM017/SM018, on genomic DNA of *S. cerevisiae* YPH500 strain) were cloned into *Nhe*I-digested yeast vector for integration into *H2* locus, harboring yest selction auxotrophic marker URA3^20^. To construct plasmid pSM13, expression cast for LacI and NLS-JUB1-X_C191_-GAL4AD synTF (primers SM019/SM009, on pSM02), six copies of JUB1 binding sites upstream of the *CYC1* minimal promoter (primers SM010/SM011, on pSM10), oligo fragment encoding *APSK* CDS, codon optimized for yeast (synthesized by GenScript, Netherlands) and *TDH1* terminator (primers SM020/SM021, on genomic DNA of *S. cerevisiae* YPH500 strain) were cloned into *Nhe*I-digested yeast vector for integration into *H5* locus, harboring yeast selection marker kanMX^20^. Integration of the pathway genes into *H1, H2*, and *H5* loci was performed by transforming donor fragments of pSM11-13 (primers SM022/SM023, SM024/SM025, and SM026/SM027, respectively), aided by Cas9-containing pCRCT plasmids, which also harbor gRNAs targeting the respective loci to produce strains yJUB1-X025, yJUB1-X026, and yJUB1-X_PAPS_ sequentially. Correct genomic integration and sequence integrity of all reporter constructs were verified by colony PCR and Sanger sequencing.

### PAPS production, qPCR, and data analysis

A yeast strain carrying the PAPS pathway was grown under IPTG-inducing conditions and used to quantify relative transcript levels as a proxy for transcriptional activity by RT-qPCR. Single colonies were grown overnight (∼16 h) in selective SC medium at 30°C with shaking at 220 rpm. Main cultures were then started at an initial OD_600_ of ∼0.1 in YPDA medium. Inducing cultures contained 2% (w/v) galactose and 20 mM IPTG^18^, while non-inducing cultures contained 2% (w/v) glucose. Cultures were grown for 4–6 h at 30°C with 220 rpm. Cells were harvested for RNA extraction (Qiagen RNeasy Kit) and DNase treatment (TURBO DNA-free Kit). cDNA was prepared by reverse transcription on the RNA (NEB LunaScript RT SuperMix Kit), and qPCR analysis was carried out (BIORAD SsoFast EvaGreen supermix with low ROX) using SYBR green with FBA1 as an internal reference gene. Relative expression levels were calculated using the ΔΔC_t_ method.

### HPLC

Metabolite extraction was carried out using the final strain. 10 ml of yeast culture was harvested by centrifugation at 1,000 g at −20 °C for 5 min. Cell pellets were washed once with 5 ml of pre-chilled 60% (v/v) methanol (−20 °C) and centrifuged again under the same conditions. Pellets were resuspended in 1 ml of pre-chilled 80% (v/v) methanol (−80 °C) and transferred to 2 ml bead-beating tubes containing 0.5 mm beads. Cells were disrupted using a homogenizer (8,000 rpm; 3 × 30 s cycles with 90 s rest), followed by incubation at −80 °C for 20 min and a second homogenization step (8,000 rpm; 2 × 20 s cycles with 90 s rest). Lysates were centrifuged at 1,000 g at −20 °C for 30 s, and the supernatant was filtered through a 0.22 μm syringe filter prior to analysis.

For quantification, a calibration curve for PAPS was generated using authentic standards. Briefly, 1 mg of PAPS was dissolved in 500 μL of methanol (MeOH) to prepare a stock solution. From this stock, 10 μL (20 μg PAPS) was diluted with 450 μL MeOH to make a working solution with 20 μg PAPS in 460 μL. Aliquots of 1–20 μL of this solution, corresponding to 0.043–0.87 μg PAPS injected, were analyzed by HPLC, and the resulting peak areas were plotted against the amount of PAPS injected to generate the calibration curve. For samples, 5 μL of each extract was injected directly, and the PAPS concentration was determined by comparing the detected PAPS peak area to the calibration curve. Mass spectrometric analyses were carried out using an Agilent 6120 UPLC–MS system equipped with a single-quadrupole mass analyzer and an electrospray ionization (ESI) source operating in both positive and negative ionization modes. Chromatographic separation was achieved on a Zorbax Eclipse Plus C18 column (2.1 × 50 mm, 1.8 μm particle size) using a UPLC pump at a flow rate of 0.8 mL min^−1^. A ternary mobile phase system consisting of methanol, water, and formic acid was employed, with mobile phase A comprising 99.9% MeOH and 0.1% HCOOH (v/v).

### Growth assays

Growth assays were performed as described for the induction experiments, with the following modifications. Galactose was used as the carbon source in a non-induction medium, and the induction medium contained both galactose and IPTG. Experimental cultures were inoculated to an initial OD_600_ of ∼0.2 in a total volume of 2 mL and grown at 30 °C in a 24-well microplate reader. OD_600_ measurements were recorded over a 34h period.

## RESULTS AND DISCUSSION

### Design of JUB1-X synTFs

A core objective of our work is to develop tunable regulators that enable genetic reprogramming of *S. cerevisiae* to balance the expression of multiple enzymatic pathways. IPTG-inducible plant-derived synTFs incorporate DBDs originating from plant TF and ADs compatible with yeast, inserted downstream of a nuclear localization signal (NLS; from the SV40 large T antigen)^18^. Among them, synTFs derived from JUB1 TF showed high inducible activation with low basal activity and tunability across synthetic promoters. Here, to further enhance their tunability, short tripeptides of valine, tyrosine, and glycine (YVG) and glycine, glutamate, and arginine (GER) were selected to minimally perturb the JUB1 NAC fold while increasing local conformational flexibility (**Fig. 1a**). This small sequence introduces flexible loops. Conserved cysteine residues were chosen for insertion of the motif because they are structurally sensitive regions within the NAC domain of NAC TFs of plants, where local rigidity can constrain functional engagement without compromising global folding. For example, Cys114 is located near the C/D subdomains and contributes to DNA binding, whereas Cys191 lies near the C-terminal boundary of the NAC domain, adjacent to the transition to more flexible regions. Targeting these positions allows selective modulation of DNA-binding adaptability and domain coupling, resulting in enhanced transcriptional output and expanded tunability while preserving the integrity of the resulting JUB1-X synTFs and cellular fitness. Upon characterization of the generated JUB1-X synTF harboring yeast GAL4 activation domain, we observed an enhanced inducibility and an expanded dynamic range across synthetic promoters harboring its binding site in one to multiple copies, compared to the parental JUB1 synTF (**Fig. 1b**). Notably, insertion of the YVG and GER motifs at selected cysteine positions increased maximal activation. Further characterization of JUB1-X variants harboring an additional yeast-compatible activation domain derived from plants, the EDLL activation domain, expands the graded promoter-dependent transcriptional responses (**Fig. 1c**).

**Figure 1.**
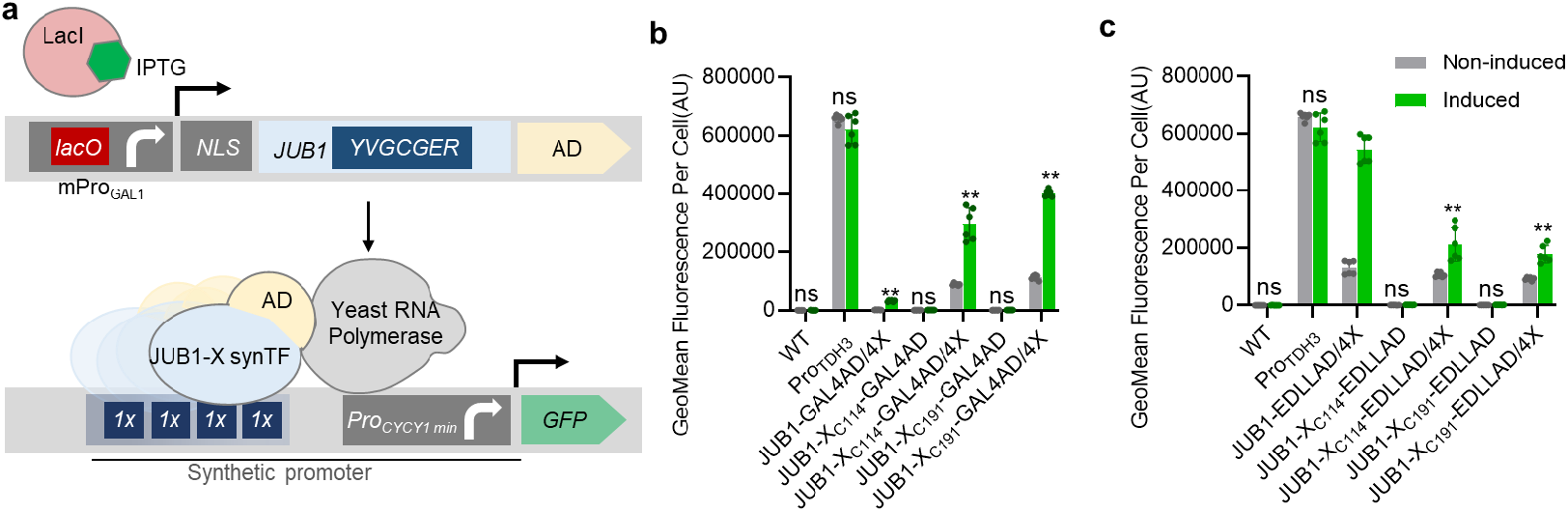
JUB1-X synTF. **a** A schematic demonstrating JUB1-X synTF established in this study. IPTG-inducible synTFs incorporate DBDs originating from the JUB1 TF of the plant, fused to activation domains compatible with yeast ^21^. within the DBD of JUB1 to add the so-called X motif to JUB1 DBD. To evaluate the performance of these synTFs in inducing gene expression, chromosomally encoded JUB1-X synTFs were placed under the control of an IPTG-responsive promoter. In the presence of IPTG, IPTG binds to LacI, forming the IPTG-LacI complex that prevents binding to the *lacO* site in the promoter, thereby activating synTF expression. Next, the nucleus-guided synTF targets its binding site (BS), located in one or more copies upstream of a minimal CYC1 promoter, thereby recruiting RNAP to the promoter. As a result, yeGFP is expressed. Transcriptional output of JUB1-X synTF harboring X motif at **b** 114 cystein and **c** 191 cystein. The CDSs of JUB1 harboring the X motif in cysteine sites 114 and 191 were inserted between the N-terminus NLS and GAL4 or EDLL ADs. The generated JUB1-X synTFs were tested for their strength to activate yEGFP reporter gene expression from the *CYC1* minimal promoter harboring four (4×) copies of the cognate binding sites. The effector and reporter constructs were chromosomally integrated into the yeast genome. The yEGFP output signal was tested both in the absence and in the presence of the inducer (IPTG). Gray, non-induction medium; green, induction medium. WT, yeast YPH500 wild type control; *Pro*_*TDH3*_, yeGFP under the control of yeast constitutive promoter *TDH3*. Mean fluorescence intensity per cell is given. Data are geometric means ± SD of fluorescence intensity from two cultures, each derived from two independent yeast colonies and measured in three technical replicates. Asterisks indicate statistically significant difference from noninduction medium (Student’s t-test; (∗) p < 0.05; (∗∗) p < 0.01; ns, non-significant). Abbreviations: AD, activation domain; AU, arbitrary units; BS, binding site; IPTG, isopropyl β-D-1-thiogalactopyranoside; JUB1, JUNGBRUNNEN1 transcription factor; *lacO*, lactose operator; LacI, lactose operon repressor; NLS, nuclear localization signal; Pro, promoter; RNAP, RNA polymerase; JUB1-derived synthetic transcription factor harboring Short tripeptide YVG and GER were placed in the N- and C-terminus of cycteins; yeGFP, yeast-enhanced green fluorescent protein. Full data are shown in **Supplementary Data S1**.

### JUB1-X synTF collection for tunable gene expression in yeast

The four JUB1-X synTFs JUB1-X_114_-GAL4AD, JUB1-X_191_-GAL4AD, JUB1-X_114_-EDLLAD, and JUB1-X_191_-EDLLAD, in combination with one or multiple copies of the JUB1 binding site, were next characterized for their ability to drive gene expression in yeast. As shown in **Figure 1**, the GAL4AD-based JUB1-X synTFs exhibited a pronounced and graded increase in transcriptional output as the number of binding sites was raised, demonstrating a high degree of JUB1 TFs’ tunability across promoter architectures. In contrast, EDLLAD-containing JUb1-X synTFs produced higher expression levels. Furthermore, JUB1-X_191_ variants consistently outperformed the JUB1-X_114_ variants. Together, these results suggest that JUB1-X synTFs are highly tunable transcriptional activators, a highly desirable feature for minimizing metabolic burden in dynamic metabolic engineering applications.

To assess the impact of JUB1-X synTF activity on growth, we cultivated strains expressing weak, medium, or strong JUB1-X synTFs under inducing conditions. As shown in **Fig. 2e**, activation of the strongest synTF, which drives the highest reporter expression, resulted in only a slight reduction in growth (green line) relative to wild-type cells (grey line). Importantly, in the absence of induction, JUB1-X synTF-expressing strains displayed growth similar to wild type (**Fig. 2f**). By contrast, constitutive GFP expression caused a clear growth defect even under non-induced conditions (blue line). Altogether, these results showed that inducible JUB1-X synTFs impose a lower metabolic burden than constitutive expression systems, enabling robust biomass accumulation before the production phase begins.

**Figure 2.**
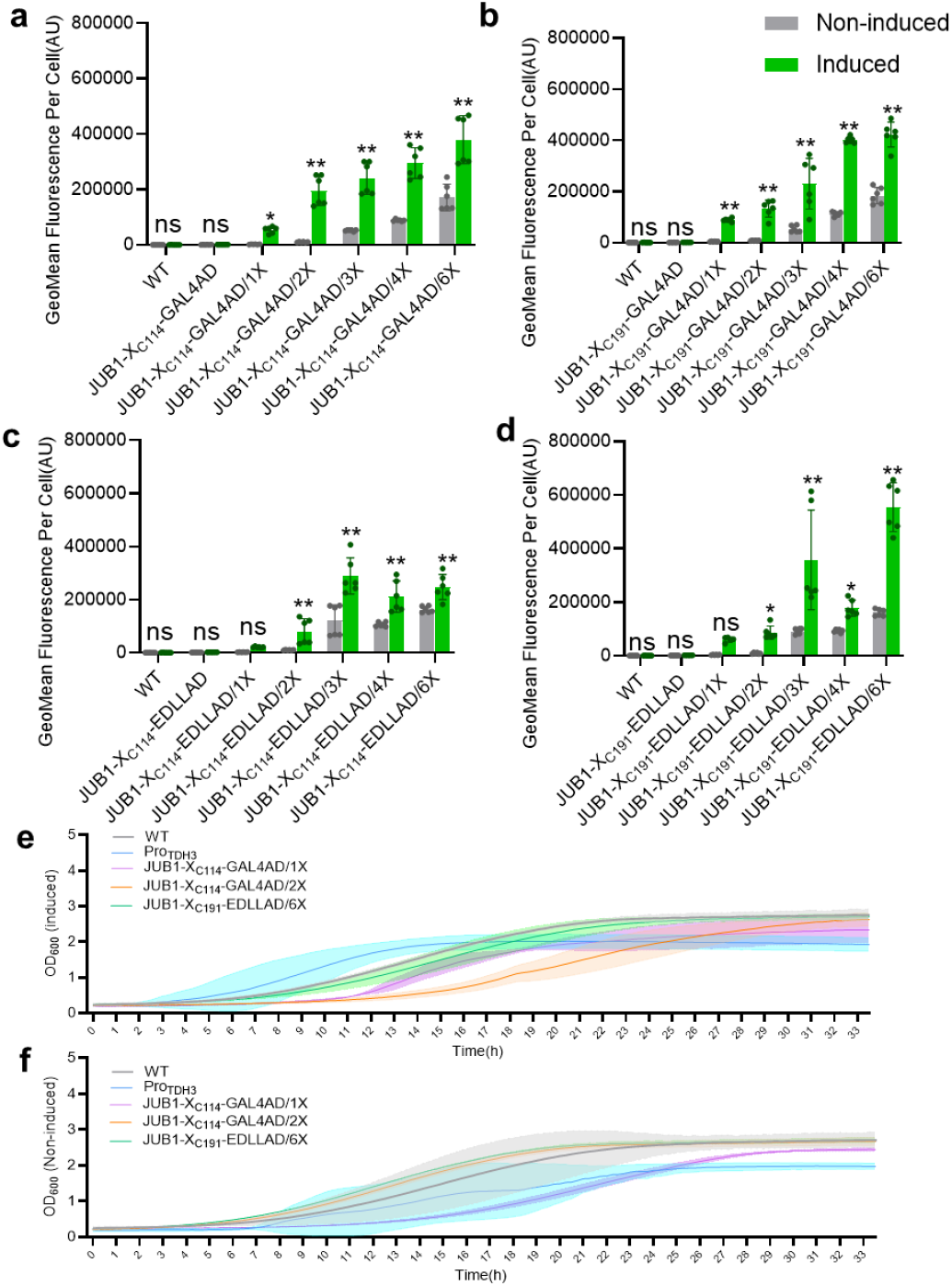
Tunable transcriptional activation by JUB1-X synthetic transcription factors through binding-site copy number modulation. Transcriptional output of four JUB1-X synTF variants, JUB1-X_114_-GAL4AD (**a**), JUB1-X_191_-GAL4AD (**b**), JUB1-X_114_-EDLLAD (**c**), and JUB1-X_191_-EDLLAD (**d**), controlling reporter gene expression in yeast. JUB1-X synTFs were expressed from an IPTG-inducible promoter and targeted synthetic promoters harboring one (1×), two (2×), or four (4×) copies of the cognate binding site upstream of a minimal *CYC1* promoter driving yEGFP expression. Both expression and reporter constructs were chromosomally integrated into the yeast genome. yEGFP fluorescence was measured in the absence (gray) and presence (green) of IPTG. Data are presented as geometric mean fluorescence intensity per cell ± SD from two independent yeast colonies, each measured in three technical replicates. Asterisks indicate statistically significant differences relative to non-induction conditions (Student’s t-test; (∗) p < 0.05; (∗∗) p < 0.01; ns, non-significant). Time course growth assay of yeast cells containing chromosomally integrated JUB1-X synTFs controlling reporter gene expression in **e** induction medium and **f** non-induction medium. The strains containing the integrated weak and medium by combining synTF JUB1-X_C114_-GAL4AD, respectively, with one and two copies of JUB1 binding sites and strong synTF by combining JUB1-X_C191_-EDLL4AD with six copies of JUB1 binding sites controlling yEGFP expression were inoculated into non-induction medium (without IPTG), and induction (with IPTG), and OD_600_ was measured over 34h. Data are presented as the mean of OD_600_ ± SD from three independent yeast colonies. Asterisks indicate statistically significant differences relative to noninduction conditions (Student’s t-test; (∗) p < 0.05; (∗∗) p < 0.01; ns, non-significant). Abbreviations: AD, activation domain; AU, arbitrary units; BS, binding site; IPTG, isopropyl β-D-1-thiogalactopyranoside; JUB1, JUNGBRUNNEN1 transcription factor; JUB1-X synTF, JUB1-derived synthetic transcription factor harboring Short tripeptide YVG and GER were placed in the N- and C-terminus of cycteins; yEGFP, yeast-enhanced green fluorescent protein. Full data are shown in **Supplementary Data S2**.

### JUB1-X synTF-derived PAPS-producing yeast strain

PAPS is synthesized from ATP and inorganic sulfate via the sequential action of ATPS and APSK, directly coupling sulfate activation to cellular energy and sulfur metabolism (**Fig. 3a**). To establish a yeast platform capable of growth-decoupled PAPS production, we first rationally selected highly efficient enzymes for PAPS biosynthesis and ATP regeneration. Previous studies have shown that ATPS from *S. cerevisiae* exhibits higher catalytic activity, whereas APSK from *E. coli* displays superior efficiency, achieving substantially higher ATP-to-PAPS conversion than fungal APSKs^21^. Based on these findings, we selected ATPS from *S. cerevisiae* and APSK from *E. coli* as the core enzymes for PAPS biosynthesis in our engineered yeast strain. Because PAPS synthesis consumes ATP and generates ADP, sustained pathway activity requires efficient ATP regeneration to prevent energetic limitation. To address this constraint, we incorporated a polyphosphate kinase (PPK2_CpaN_) from *Pseudomonas* sp., which catalyzes ATP regeneration from ADP using inorganic polyphosphate as a phosphate donor^21^. This strategy has been shown to markedly enhance ATP availability and overall PAPS conversion efficiency in vitro, and is compatible with yeast physiology, as *S. cerevisiae* tolerates polyphosphate supplementation. The genes encoding PPK2_CpaN_, ATPS, and APSK were placed under the control of synTFs medium-to-strong JUB1-X_C114_-GAL4AD/2X, JUB1-X_C191_-GAL4AD/4X, and JUB1-X_C191_-GAL4AD/6X, respectively, and were chromosomally integrated into the yeast genome to generate strain yJUB1-X_PAPS_. Upon IPTG induction, transcriptional levels of all three pathway genes increased significantly compared to noninduced conditions, reaching 204-, 170-, and 54-fold induction under the control of JUB1-X_C114_-GAL4AD/2X, JUB1-X_C191_-GAL4AD/4X, and JUB1-X_C191_-GAL4AD/6X, respectively (**Fig. 3b**). Correspondingly, expression levels were 3.3-, 9.7-, and 12.5-fold higher than those of housekeeping genes. Consistent with the observed induction of pathway gene expression, IPTG induction led to robust intracellular accumulation of PAPS in strain yJUB1-X_PAPS_ (**Fig. 3b**). Under simple batch flask culture conditions, the engineered strain resulted in PAPS accumulation of 21.4 ± 5.8 mg g^−1^ cdw (**Fig. 3c**) only after only 5 hours of inducing the expression of JUB1-X synTFs controlling expression of PAPS enzymes, while no detectable PAPS was obtained for non-induced condition and wild-type strain (see also **Supplementary Fig. S1**). Importantly, induction of the PAPS pathway after only 5 hours of growth (the same time as the time to characterize JUB1-X synTFs, **Fig. 1** and **Fig. 2**), which corresponds to the mid-exponential growth phase, ensures that cells remain highly metabolically active while having accumulated sufficient biomass. These results confirm the high transcriptional capacity of JUB1-X synTFs for metabolic engineering applications in yeast *S. cerevisiae*. To evaluate whether pathway induction imposes an additional growth burden, we next monitored the growth dynamics of strains expressing the APSK–ATPS–PPK2 genes under synTF control. As shown in **Fig. 3d**, induction of the pathway resulted in only modest alterations to growth kinetics compared with the wild-type controls. Interestingly, the PAPS-producing strain showed reduced growth under non-induced conditions but partial recovery upon induction, while the wild-type strain showed similar growth in both induced and non-induced media. This suggests that leaky expression of the heterologous pathway (under non-induced conditions) led to an imbalance in metabolic flux, possibly the accumulation of intermediates, and thereby, a linked growth burden. Such effects have been previously recognized in metabolic engineering, where optimizing expression levels to balance pathway flux can minimize growth inhibition and the buildup of toxic intermediates^22^. Together, these results establish a modular and energetically balanced PAPS-producing yeast strain in which enzyme expression strength and timing can be precisely controlled using next-generation JUB1-X synTFs.

**Figure 3.**
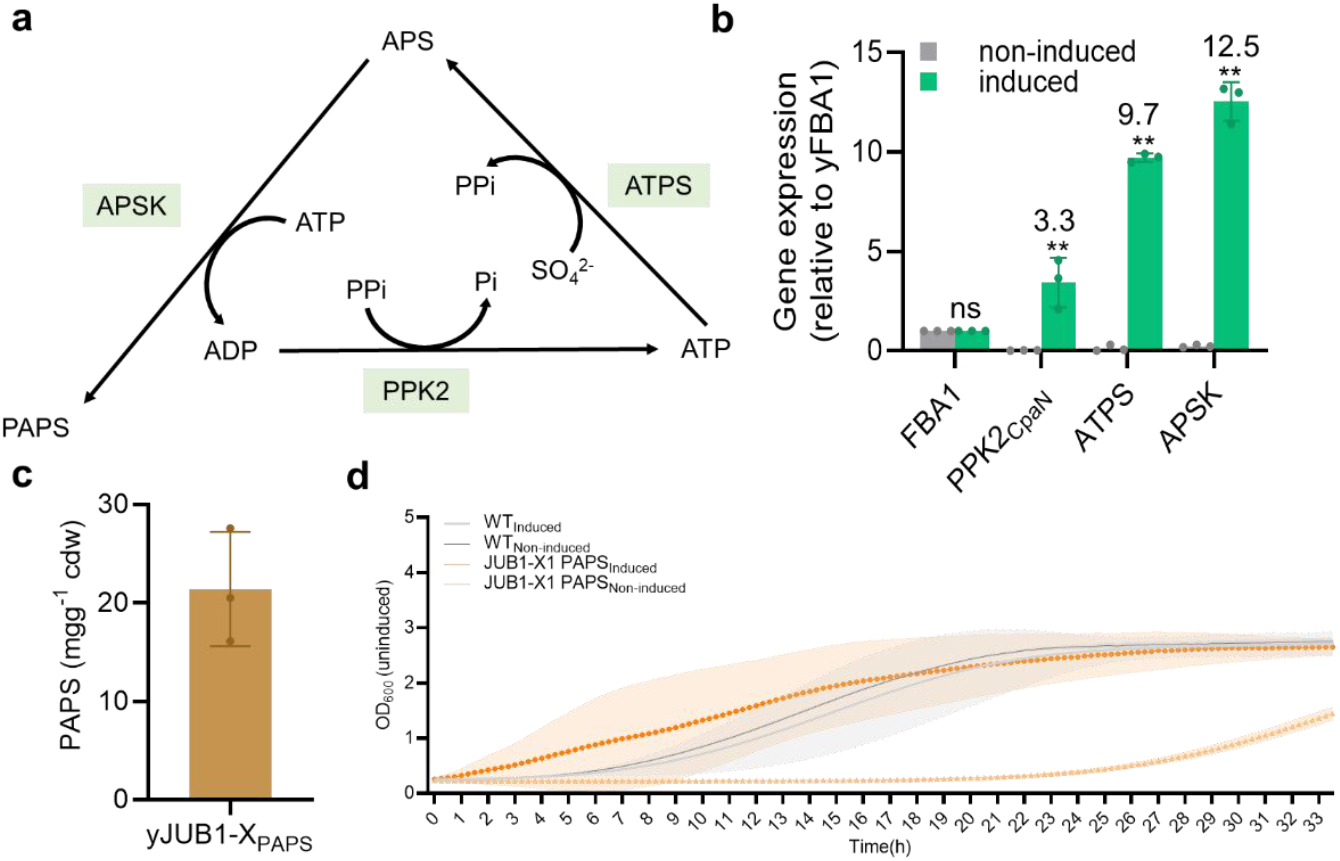
Engineering platform for improved PAPS production in yeast. **a** Schematic illustration of PAPS production. To redirect yeast metabolism toward PAPS production, overexpression of APSK, ATPS, and PPK2 (green) is needed. APSK and ATPS convert ATP to PAPS, producing ADP. PPK2 utilizes polyphosphate as a phosphate donor to regenerate ATP from ADP. **b** Relative transcriptional level of PAPS-encoding enzymes using JUB1-X synTFs in yeast. The inducible JUB1-X synTFs JUB1-X_C114_-GAL4AD/2X, JUB1-X_C191_-GAL4AD/4X, and JUB1-X_C191_-GAL4AD/6X were used to express enzymes PPK2_CpaN_, APSK, and ATPS, respectively, to generate strain yJUB1-X_PAPS_. Transcription level of pathway genes was measured in the absence (gray) and presence (green) of IPTG after 5h. **c** PAPS production in strain yJUB1-X_PAPS_. Production of PAPS in the generated strain yJUB1-X_PAPS_ was measured in the presence (green) of IPTG after 5h at OD_600_ ∼ 2.0 using HPLC. No detectable PAPS was observed in the non-induction medium or in YPH500 (data not shown). **d** Time course growth assay of yeast cells producing PAPS under the control of JUB1-X synTFs. The strain yJUB1-X_PAPS_ was inoculated into both induction and non-induction media, and OD600 was measured over 34 h. Data are presented as the mean OD_600_ ± SD from three technical replicates. Asterisks indicate statistically significant differences relative to the non-induction condition (Student’s t-test; (∗) p < 0.05; (∗∗) p < 0.01; ns, non-significant). Abbreviations: ATP, adenosine 5′-triphosphate. ADP, adenosine 5′-diphosphate; PAPS, 3′-phosphoadenosine-5′-phosphosulfate; ATPS, ATP sulfurylase; APSK, APS kinase; PPi, inorganic pyrophosphatase; PPK, polyphosphate kinase; IPTG, isopropyl β-D-1-thiogalactopyranoside; JUB1, plant JUNGBRUNNEN1 transcription factor; JUNGBRUNNEN1 transcription factor; JUB1-X synTF, JUB1-derived synthetic transcription factor harboring short tripeptide YVG and GER were placed in the N- and C-terminus of cycteins; GAL4AD, yeast GAL4 activation domain; ‘2X’, ‘4X’, and ‘6X’ indicate two, four, and six copies of the synTF binding site, respectively; HPLC, high performance liquid chromatography; cdw, cell dry weight.

## Discussion

Precise control of pathway expression strength and timing is essential for efficient metabolic engineering in *S. cerevisiae*, particularly for energetically demanding biosynthetic processes. In this study, we developed next-generation JUB1-X synTFs that expand the tunable expression range of plant-derived synTFs while maintaining minimal impact on cellular fitness. We showed that rational modification of conserved regions within the JUB1 NAC DNA-binding domain yields a graded set of inducible regulators compared to previously reported JUB1-based synTFs^18^.

A major bottleneck in sulfation pathways is the limited intracellular availability of PAPS, which is biosynthesized, tightly regulated, and energetically costly. To avoid relying on constitutive overexpression or complex cofactor-recycling systems^15^, we implemented JUB1-X synTFs to express PAPS biosynthetic enzymes. By inducing pathway expression only at the appropriate time and after the biomass accumulation phase, the engineered strain yJUB1-XPAPS produced 21.4 ± 5.8 mg g^−1^ cdw PAPS in a simple batch flask culture, whereas no PAPS was detected under noninduced conditions.

Our system requires only the addition of IPTG to the culture medium, offering a substantially simple and scalable platform, compared to other available engineered available system^15^. Future efforts could focus on further tuning induction timing, modifying the copy number of JUB1-X synTF, or integrating dynamic feedback systems to maximize yields. Importantly, the engineering principle established here, enhancing tunability through modification of conserved regions within plant-derived DBD, is not limited to JUB1. It is broadly applicable to other synTFs, including those described by Naseri *et al*^18^, which employ diverse plant DNA-binding domains. The modularity and tunability of JUB1-X synTFs make them promising tools for production in a wide range of industrially relevant pathways. This strategy could be extended to other cofactor-dependent pathways where production competes with growth, such as NADPH-dependent polyketide or flavonoid biosynthesis, SAM-dependent methylation reactions, and acetyl-CoA-intensive terpene or lipid pathways. In summary, this work introduces JUB1-X synTFs as an improved, tunable regulatory tool for dynamic metabolic engineering in yeast. By enabling growth-decoupled control of multigene pathways, JUB1-X synTFs support the efficient production of energetically demanding cofactors, such as PAPS. This regulatory strategy is broadly applicable to improve the transcriptional activity of other synTFs derived from dCas9 or other heterologous TFs, and to optimize the engineering of other complex biosynthetic pathways that require synchronized and adjustable gene expression in *S. cerevisiae* and other microbial cell factories.

## Abbreviations Used

1×, 2×, 4×, 6×: Number of copies of a synTF binding site
AD: Activation domain
ADS: Amorphadiene synthase
APSK: Adenosine 5′-phosphosulfate kinase
ATPS: ATP sulfurylase
AU: Arbitrary units
BS: Binding site
CDW: Cell dry weight
Cys: Cysteine
DBD: DNA-binding domain
EDLLAD: EDLL activation domain
GAL4AD: Yeast GAL4 activation domain
GER: Tripeptide glycine–glutamate–arginine
IPTG: Isopropyl β-D-1-thiogalactopyranoside
JUB1: JUNGBRUNNEN1 transcription factor
JUB1-X synTF: JUB1-derived synthetic transcription factor with short tripeptides YVG and GER inserted at N- and C-terminal cysteine positions
JUB1-X114: JUB1-X synTF with tripeptides at cysteine 114
JUB1-X191: JUB1-X synTF with tripeptides at cysteine 191
LacI: Lactose operon repressor
lacO: Lactose operator
NLS: Nuclear localization signal
PAPS: 3′-Phosphoadenosine 5′-phosphosulfate
PPA: Inorganic pyrophosphatase
PPi: Inorganic pyrophosphate
PPK: Polyphosphate kinase
Pro: Promoter
ProTDH3: Yeast constitutive TDH3 promoter
RNAP: RNA polymerase
WT: Wild type
yEGFP: Yeast-enhanced green fluorescent protein
yJUB1-XPAPS: Yeast strain expressing PAPS pathway USINF JUB1-X synTFs
YVG: Tripeptide tyrosine–valine–glycine

## ASSOCIATED CONTENT

### Supporting Information

The authors declare that the data supporting the findings of this study are available within the paper and its Supplementary Information files. Should any raw data files be needed in another format, they are available from the corresponding author upon reasonable request.

## AUTHOR INFORMATION

## Authors

**Madhushruti Borah**. - Institut of Biology, Humboldt-Universität zu Berlin, Philippstrasse 13, 10115 Berlin, Germany;

**Shanna Gu**. - Institut of Biology, Humboldt-Universität zu Berlin, Philippstrasse 13, 10115 Berlin, Germany;

**Essa M. Saied** - Institute of Chemistry, Humboldt-Universität zu Berlin, Brook-Taylor-Str. 2, 12489 Berlin, Germany; Chemistry Department, Faculty of Science, Suez Canal University, Ismailia 41522, Egypt;

**Christoph Arenz** - Institute of Chemistry, Humboldt-Universität zu Berlin, Brook-Taylor-Str. 2, 12489 Berlin, Germany;

**Mattheos Koffas** - Department of Chemical and Biological Engineering, Rensselaer Polytechnic Institute, Troy, NY 12180, USA;

## Author contributions

GN conceived and supervised the project. SG designed synthetic transcription factors and performed their characterization in yeast. MK designed the PAPS engineering strategy. MB designed, constructed, and produced the PAPS-producing yeast strain. The experiments and data analysis were carried out by MB and SG. Chromatography experiments were performed and analyzed by EMS and CA. The manuscript was written by GN, with contributions from SG and MB. All authors take full responsibility for the content of the paper.

## Notes

The authors declare no competing interests.

## Acknowledgements

Gita Naseri would like to acknowledge support from the Institute for Biology, Humboldt-Universität zu Berlin, and Neel H. Shah (Columbia University, New York, NY, USA) for his guidance in designing synthetic transcription factors.

## Supporting Information description

**Table S1**. Sequence of plasmids generated in the present study.

**Table S2**. Sequence of primers used in the present study.

**Supplementary Data S1**. JUB1-X synTF.

**Supplementary Data S2**. Tunable transcriptional activation by JUB1-X synthetic transcription factors through binding-site copy number modulation

**Supplementary Data S3**. Engineering platform for improved PAPS production in yeast.

---

**Supplementary Figure S1.**
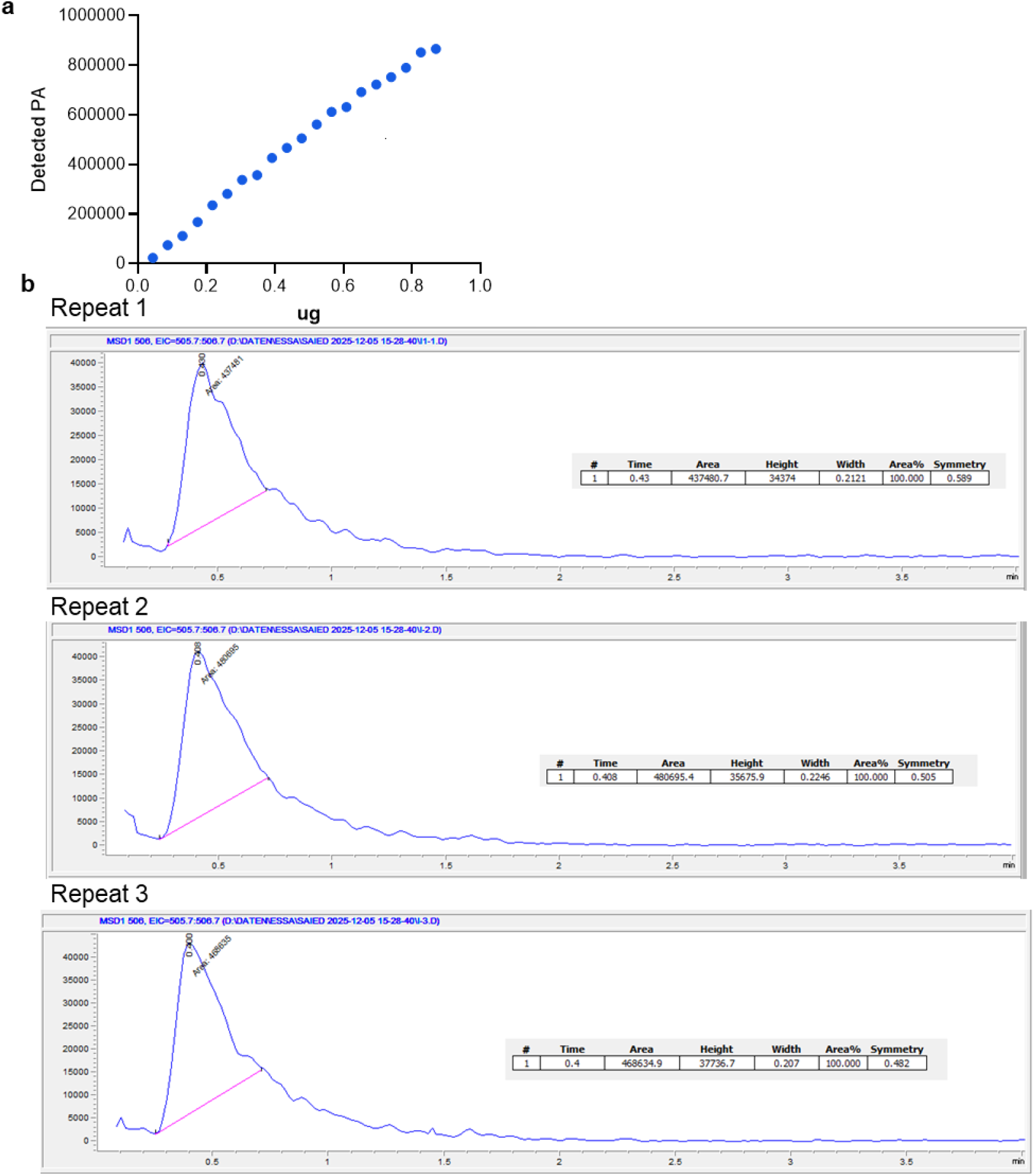
Method for HPLC analysis of PAPS-producing samples. **a** PAPS calibration curve. **b** PAPS chromatogram for strain JUB1-X_PAPS_. Abbreviations: PAPS, 3’-phosphoadenosine 5’-phosphosulfate.

